# Molecular architecture of the *C. elegans* centriole

**DOI:** 10.1101/2022.05.09.491129

**Authors:** Alexander Woglar, Marie Pierron, Fabian Zacharias Schneider, Keshav Jha, Coralie Busso, Pierre Gönczy

## Abstract

Uncovering organizing principles of organelle assembly is a fundamental pursuit in the life sciences. *C. elegans* was key in identifying evolutionary conserved components governing assembly of the centriole organelle. However, localizing these components with high precision has been hampered by the minute size of the worm centriole, thus impeding understanding of underlying assembly mechanisms. Here, we used Ultrastructure Expansion coupled with STimulated Emission Depletion microscopy (U-Ex-STED), as well as electron microscopy (EM) and tomography (ET), to decipher the molecular architecture of the worm centriole. Achieving an effective lateral resolution of ∼14 nm, we localize centriolar and PeriCentriolar Material (PCM) components in a comprehensive manner with utmost spatial precision. We uncovered that the procentriole assembles from a location on the centriole margin characterized by SPD-2 and ZYG-1 accumulation. Moreover, we found that SAS-6 and SAS-5 are present in the nascent procentriole, with SAS-4 and microtubules recruited thereafter. We registered U-Ex-STED and EM data using the radial array of microtubules, thus allowing us to map each centriolar and PCM protein to a specific ultrastructural compartment. Importantly, we discovered that SAS-6 and SAS-4 exhibit a radial symmetry that is offset relative to microtubules, leading to a chiral centriole ensemble. Furthermore, we establish that the centriole is surrounded by a region from which ribosomes are excluded and to which SAS-7 localizes. Overall, our work uncovers the molecular architecture of the *C. elegans* centriole in unprecedented detail and establishes a comprehensive framework for understanding mechanisms of organelle biogenesis and function.

## Introduction

Centrioles are membrane-less organelles that were present in the last common ancestor of eukaryotes (reviewed in 1). In cells with flagella or cilia, centrioles act as basal bodies that template the formation of these structures. Moreover, in animal cells, centrioles form the core of the centrosome, which organizes microtubules and is thereby critical for fundamental cellular processes, including polarity and division (reviewed in 2). In most organisms, centrioles are cylindrical organelles ∼500 nm high x ∼250 nm wide, with a 9-fold radially symmetric distribution of microtubules (reviewed in 3, 4). These centriolar microtubules are organized in triplets in the proximal region of the organelle and in doublets in its distal region. Triplet and doublet microtubules are twisted in a clockwise direction with respect to the microtubules when viewed from the distal end of the centriole, resulting in the characteristic chiral geometry of the organelle. This 9-fold radially symmetric architecture is also imparted onto the ciliary and flagellar axoneme that stem from centriolar microtubules, and might be evolutionarily conserved because it provides an optimal geometry for axonemal motility. Despite important progress in recent years, the detailed molecular architecture of the centriole, including the root of its characteristic chirality, remains incompletely understood.

There are variations in the architectural features of centrioles in some systems, which are usually correlated with the absence or reduction of ciliary and flagellar motility (reviewed in 5). For instance, in the nematode *Caenorhabditis elegans*, motile cilia and flagella are absent, and the sperm moves in an amoeboid fashion. Perhaps in the absence of evolutionary pressure for ciliary and flagellar motility, centrioles are smaller (∼175 nm high x ∼120 nm wide in the embryo (6–8), and comprise a radial arrangement of 9 microtubule singlets instead of the usual triplets and doublets (9). Electron microscopy (EM) of centrioles in the *C. elegan*s embryo revealed ultrastructural compartments besides microtubules, including 9 peripheral paddlewheels, as well as the central tube and, more centrally still, the inner tube (6–8). EM analysis of embryonic centrioles also led to the notion that each paddlewheel is offset with respect to its accompanying microtubule, with a clockwise twist when viewed from the distal end, resulting in a chiral ensemble (8). Whether chirality of the *C. elegans* centriole is apparent more centrally in the organelle, where the assembly process is thought to initiate, is not known.

As in other systems, starting approximately at the onset of S phase, the two resident centrioles in *C. elegans* each seed the assembly of a procentriole in their vicinity, such that four centriolar units are present during mitosis, two per spindle pole. Comprehensive genetic and functional genomic screens conducted in *C. elegans* led to the discovery of six components essential for procentriole formation, which are largely conserved in overall structure and function across eukaryotic evolution (reviewed in 10–12). Molecular epistasis experiments uncovered the order in which proteins essential for procentriole formation are recruited to the worm organelle (7, 13). These experiments established that SAS-7 and SPD-2 (Cep192 in humans) are first recruited to the resident centriole. Thereafter, the kinase ZYG-1 (Plk4 in humans) directs the interacting coiled-coil proteins SAS-6 (HsSAS-6 in humans) and SAS-5 (STIL in humans) to the procentriole assembly site. This is followed by SAS-4 (CPAP in human) recruitment to the procentriole, a protein thought to enable the addition of microtubules to the SAS-6/SAS-5 scaffold. Relatives of SAS-7, SPD-2, ZYG-1, SAS-6, SAS-5 and SAS-4 in other systems are recruited in a similar sequence and exert analogous functions in procentriole formation (reviewed in 10–12).

SAS-6 is the main building block of a scaffold referred to as the cartwheel, which is thought to contribute to imparting the 9-fold radial symmetry of the organelle (14, 15). Whereas SAS-6 proteins in other systems self-assemble into ring-containing polymers that stack to form the cartwheel, structural and biophysical evidence obtained with the *C. elegans* protein has led to the suggestion that SAS-6 forms a steep spiral (16). However, whether this is the case *in vivo* has not been addressed.

HYLS-1 and SAS-1 are two additional *C. elegans* centriolar proteins that are dispensable for procentriole assembly. However, HYLS-1 is needed for generating non-motile cilia (17), whereas SAS-1 is critical for maintaining the integrity of the organelle once formed (18). In addition, the Polo-like kinase PLK-1 is present at centrioles in the early worm embryo (19). As in other systems, *C. elegans* centrioles recruit the core PeriCentriolar Material (PCM), thus forming the centrosome, which acts as a microtubule organizing center (reviewed in (20). Assembly of the core PCM in *C. elegans* relies on the interacting proteins SPD-2 (21, 22) and SPD-5 (23), as well as on SAS-7 (8) and PCDM-1 (24). Furthermore, the γ-tubulin protein TBG-1 (25, 26), together with the γ-tubulin interacting proteins GIP-1 and GIP-2 (26), as well as the γ-tubulin partner MZT-1 (27) are present in the core worm PCM. Additional proteins, including PLK-1 and AIR-1 (28, 29), as well as TAC-1 and ZYG-9 (30, 31), are recruited to this core PCM when the centriole matures in mitosis in the embryo, leading to increased microtubule nucleation. Despite the probably near comprehensive list of component parts of the centriole and core PCM in *C. elegans*, the very small dimensions of the worm organelle have thus far prevented localizing with precision where each component resides, thus limiting understanding of how they function.

The molecular architecture of the centrioles has been investigated using 3D-Structured Illumination Microscopy (SIM) or STED super-resolution microscopy in other systems, including human cells and Drosophila (32–34), where the organelle is larger than in *C. elegans*. Moreover, ultrastructure expansion (U-Ex) microscopy has been utilized to investigate the molecular architecture of centrioles from human cells (35, 36). In this method, the sample is embedded in a gel that is then expanded isotropically several fold, thus likewise expanding the effective resolution (37). SIM, STED and U-Ex have enabled placing in a more refined manner a subset of components in the centriole map of these systems. However, the resolution achieved with these approaches would be likely insufficient to resolve the molecular architecture of the minute worm centriole.

Here, we set out to map in a comprehensive manner and with utmost precision the distribution of centriolar as well as core PCM component in the gonad of *C. elegans*. Considering the minute size of the worm centriole, we combined U-Ex and STED, reaching an effective lateral resolution of ∼14 nm. Using mainly endogenously tagged components and validated antibodies, we could thus determine with exquisite precision the localization of twelve centriolar and core PCM proteins. Of particular interest, this revealed that SAS-6 and SAS-4 exhibit an angular offset with respect to the microtubules, resulting in a chiral arrangement in the organelle center. Moreover, we acquired a large corresponding EM data set, which we overlaid with the U-Ex-STED images to map each centriolar protein to a specific ultrastructural compartment of the organelle. Overall, we uncovered the molecular architecture of the *C. elegans* centriole and provide an unprecedented framework for a mechanistic dissection of centriole assembly and function.

## Results

### Combining nuclei spreading and U-Ex microscopy for improved resolution of centrioles

We set out to analyze the molecular architecture of the *C. elegans* centriole with utmost spatial resolution, using the adult hermaphrodite gonad as an experimental system (Fig. 1A). The distal part of the syncytial gonad (the “mitotic zone” from here on) constitutes a stem cell pool where nuclei undergo cell cycles characterized by short G1 and M phases, with merely ∼2% of nuclei being in one of these two phases combined (38). Once nuclei have traveled far enough from the distal end of the gonad, they undergo pre-meiotic S phase and enter meiotic prophase I, a prolonged G2 phase during which meiotic recombination occurs.

**Figure 1:**
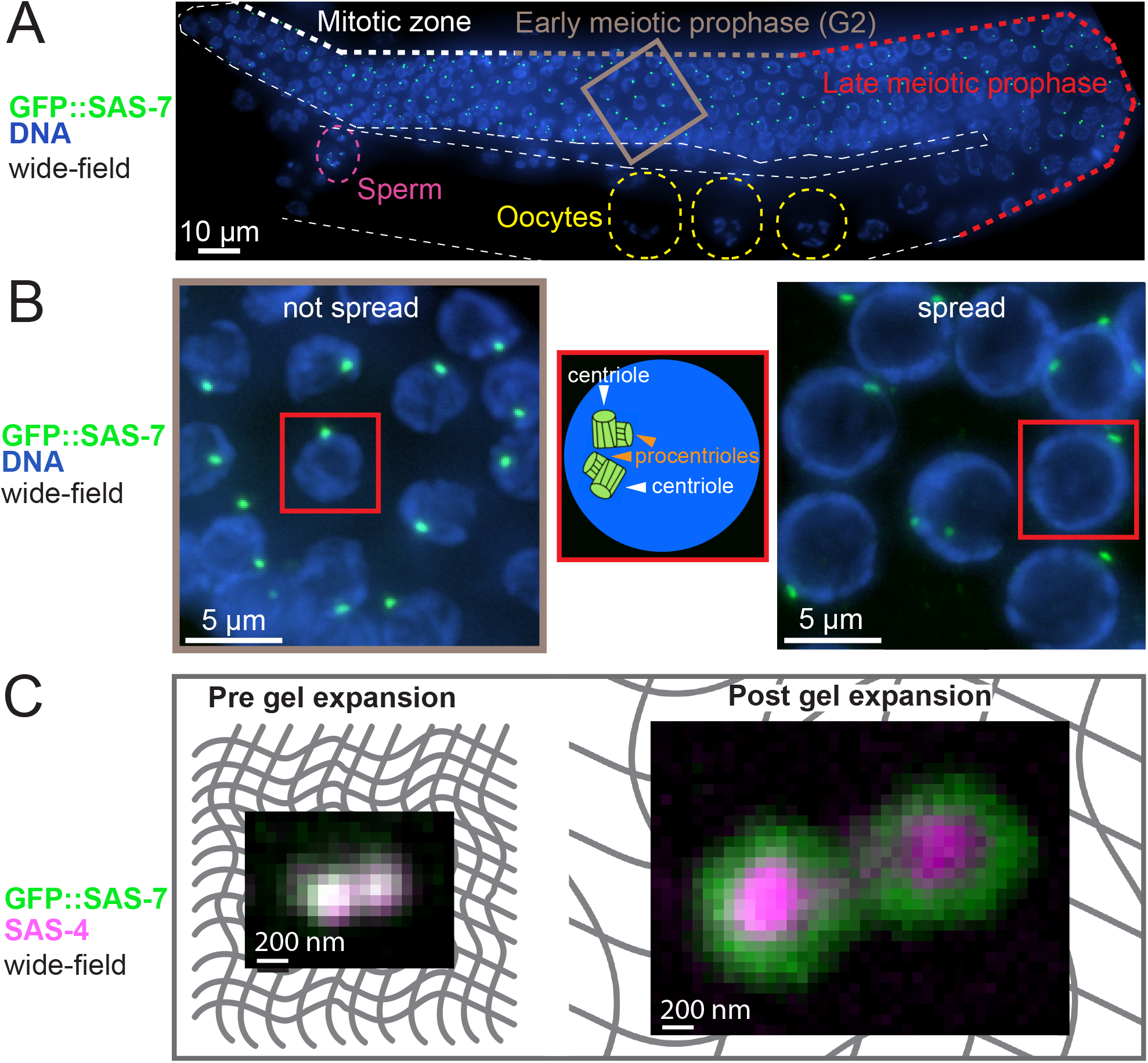
Combining gonad nuclei spreading and U-Ex microscopy to analyze worm centrioles. (A) Widefield imaging of ethanol-fixed worm expressing GFP::SAS-7. One layer of nuclei of the gonad is max intensity Z-projected (in this case a height of 6.25 µm). White, grey and red bold dashed line indicate progression through the gonad from the mitotic zone to early and then late meiotic prophase; other white dashed line outlines the gonad. Grey box is magnified in (B). Yellow dashed regions mark three oocytes, purple dashed region the spermatheca. Note that centrioles are eliminated in oocytes, prior to fertilization. (B) (Left) Magnification of grey box region from (A). (Middle) Schematic representation of a single nucleus shown in the left and right panels. (Right) Early prophase region of a spread gonad from a worm expressing GFP::SAS-7. Note that spread nuclei are flattened and thus occupy a larger area compared to not spread nuclei. (C) Widefield imaging of centrioles in the early prophase region of the gonad from worms expressing GFP::SAS-7 before (left) and after (right) gel expansion. Grey mesh in the background represents the gel matrix.

The gonad can be easily extruded from the animal and contains hundreds of nuclei, which are almost all in S or G2 phases of the cell cycle. Since procentriole formation begins in early S phase, most gonad nuclei harbor two pairs of centriole/procentriole, which are usually in close proximity to one another and cannot be resolved by Immuno-Fluorescence (IF) in widefield microscopy, usually appearing instead as a single focus (Fig. 1B, left). We took two steps to improve the spatial resolution for our analysis. First, nuclei from extracted gonads were adhered as a single layer to a coverslip using mild chromatin spreading (39), resulting in superior detection by IF since the specimen is closer to the coverslip. Moreover, the pool of cytoplasmic proteins, which normally contributes to poor signal to noise ratio that impedes detection of centriolar components, is largely washed out in this manner (Fig. 1B, right). Second, we adapted previously validated ultrastructure gel expansion methods (U-Ex) (35, 37), reaching ∼5 fold isotropic expansion of the specimens (Fig. 1C, Material and Methods). Combination of spreading with U-ExM enabled us to distinguish centriole and procentriole with widefield microscopy (Fig. S1A), as well as to localize components to distinct regions within the *C. elegans* centriole (Fig. 1C, right).

### Procentriole assembly: onset and maturation

We investigated the distribution of twelve centriolar and core PCM components. As detailed in Table S1 and the Materials and Methods section, with the exception of mCherry::HYLS-1, we visualized each protein as an endogenously N-terminally [N] or C-terminally [C] tagged component, a tagged version expressed under the endogenous promoter in the absence of the endogenous component and/or previously validated antibodies against the endogenous protein. We found that only three of the twelve components, SAS-6, SAS-5 and SAS-4, localize to both centriole and procentriole during S and G2. Using signal intensity in 3D-SIM images as a proxy for protein amount, we found that there is an undistinguishable amount in the centriole and the procentriole for both SAS-5 and SAS-6 (Fig. 2A). In contrast, the amount of SAS-4 in the procentriole is on average ∼10 times lower than it is in the centriole and also very variable (Fig. 2A).

**Figure 2:**
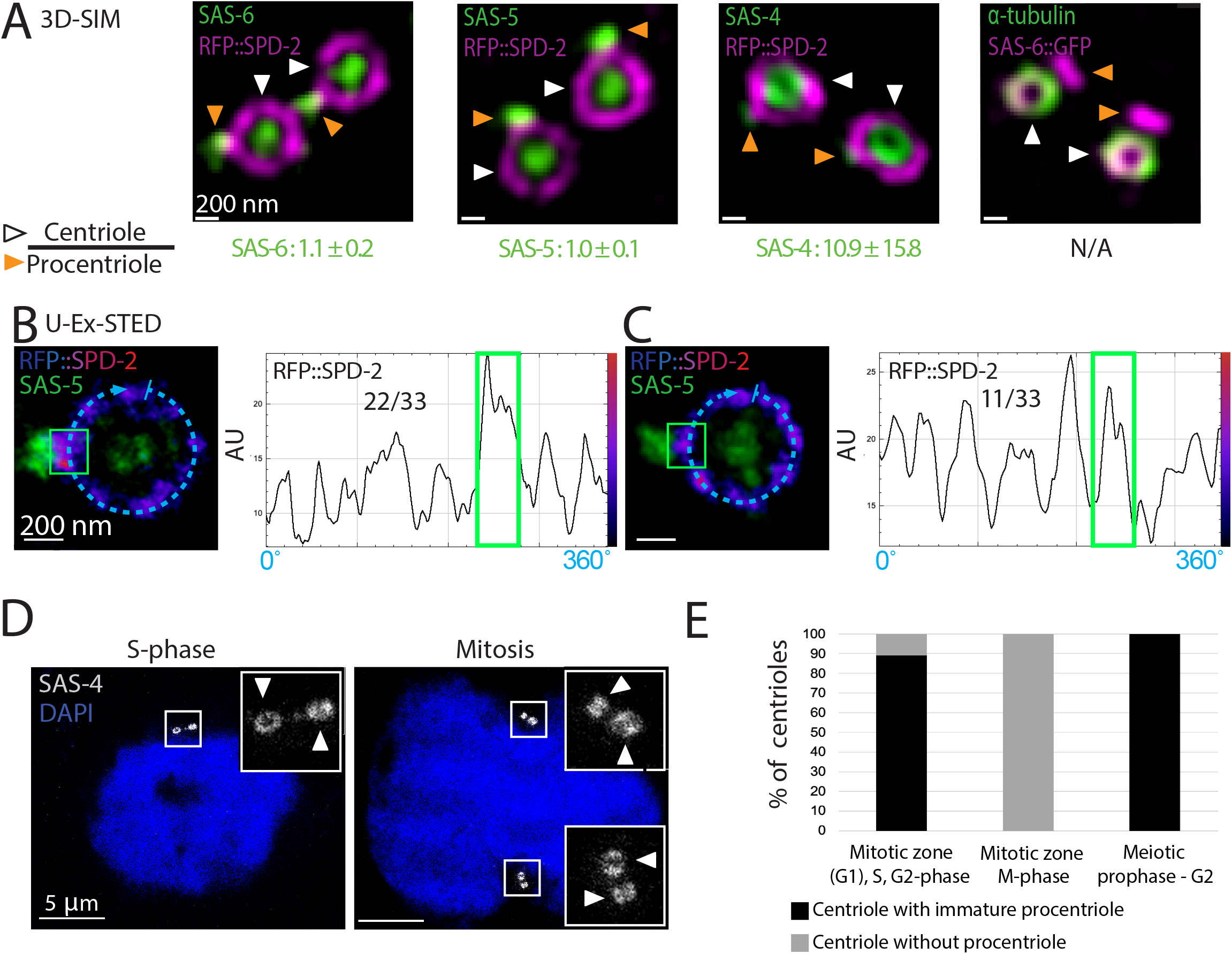
Composition and maturation of the procentriole. (A) 3D-SIM sum intensity Z-projected images of expanded centrioles from early meiotic prophase stained for the indicated proteins. White arrowheads point to centrioles, orange arrowheads to procentrioles. The presence of a ring-like distribution of RFP::SPD-2 (magenta) around one of the two green foci (SAS-6, SAS-5 or SAS-4) served to identify the centriole. The numbers below the images represent the ratio between the fluorescence of the indicated component in the centriole versus the procentriole (N= SAS-6: 8, SAS-5: 17, SAS-4: 18). Here and in all other figures, scale bars within a series represent the same length (e.g. 200 nm in this case). (B, C) (Left) U-Ex-STED of centrioles from early meiotic prophase revealing RFP::SPD-2 and SAS-5 distribution. RFP::SPD-2 signals are displayed with the LUT “Fire” (low intensities in blue, high intensities in magenta and red). (Right) Corresponding signal intensity profiles along the dashed line depicted in the image (10 pixels wide). The green boxes in the images and the graphs indicate RFP::SPD-2 located under the procentriole identified by SAS-5. In the majority of cases, the RFP::SPD-2 signal is wider and brighter below the procentriole than anywhere else in the centriole (B, 22/33), whereas such enrichment could not be detected in the remainder cases (C, 11/33). (D) U-Ex-STED of nuclei in either S-phase (left) or mitosis (right) from the mitotic zone of the gonad. White arrowheads point to centrioles fully decorated with SAS-4 (mature centrioles). (E) Percentage of centrioles (identified by SAS-6 surrounded by SPD-2) with or without a neighboring immature procentriole (identified by SAS-6 not surrounded by SPD-2) during the indicated stages. Nuclei in mitosis were identified by DNA staining as having the most condensed chromatin in the miotic region of the gonad; N= Mitotic zone (G1), S, G2-phase: 37, Mitosis (mitotic zone): 7, Early meiotic prophase: 46.

In addition to the invariable presence of SAS-6 and SAS-5 at the procentriole, we found using U-Ex-STED that ZYG-1 accumulates at the base of the procentriole in S and G2 in the mitotic region, as well as during early meiotic prophase (Fig. S1B). In contrast to SAS-6 and SAS-5, in the mitotic zone, ZYG-1 levels in the mother centrioles exhibited high variability, likely linked with cell cycle progression. We speculate that levels of ZYG-1 are higher in S phase and lower in G2 in the mitotic region because similarly low levels were observed during the prolonged G2 of meiotic prophase (Fig. S1B). Moreover, we found SPD-2 to be present in a ring around the centriole, abutting the base of the procentriole (Fig. 2B). Interestingly, SPD-2 radial distribution is not uniform, but is often enriched at the site of procentriole formation as evidenced by line scan analysis (Fig. 2B, 2C, 22/33 cases). We speculate that such an enrichment may reflect ZYG-1-mediated modification of SPD-2 to serve as a platform for procentriole formation or local increase of SPD-2 as a result of procentriole formation.

Previous analysis in the one-cell stage embryo established that the procentriole acquires SAS-4 and microtubules after SAS-6/SAS-5 recruitment (7, 13). Using U-Ex-STED, we found in the gonad that the procentriole likewise harbors little SAS-4 initially and that more protein is recruited at prometaphase, resulting in similar levels of SAS-4 in the centriole and the procentriole by then (Fig. 2D). This maturation coincides with the loading of microtubules onto the procentriole (Fig. S1C). As expected from these observations, ∼90% of centrioles harbor an immature procentriole during S and G2 phases in the mitotic region, while only centrioles without an accompanying procentriole are observed by the time of mitosis, when the procentriole disengaged and matured into a centriole (Fig. 2E). Furthermore, during the prolonged G2 of meiotic prophase that follows, all centrioles are again accompanied by an immature procentriole (Fig. 2E). These observations taken together indicate that centriole formation in the gonad is characterized by two steps: an initial rapid formation of a procentriole harboring SAS-6 and SAS-5, followed briefly before M phase by the recruitment of other components, including SAS-4 and microtubules. Interestingly, this coincides with the time during which centrioles recruit PCM (see below and reviewed in 20) and start to organize the spindle, potentially suggestive of a functional link between procentriole maturation and PCM expansion.

### U-Ex-STED reveals consecutive ring-like distribution of *C. elegans* centriolar proteins

We proceeded to comprehensively uncover the precise distribution of centriolar and core PCM proteins using U-Ex-STED. We used top views of centrioles to determine the radial distribution of component proteins (Fig. 3A). Remarkably, except for ZYG-1 (see Fig. S1B), top views of the centriole revealed that all components exhibit a ring-like distribution, with distinct diameters. To analyze the position of each component with respect to the others, we determined the diameter of each ring relative to that of α-tubulin, which was used as an invariant reference in this analysis (Fig. 3B). To verify the validity of this approach, we co-stained α-tubulin with two different antibodies, finding that the two signals colocalize and that the corresponding rings hence exhibit the same diameter (without correction for the expansion factor: C-terminus 1.44 ± 0.12 nm, N-terminus 1.43 ± 0.11 nm, p=0.81, N=19; Fig. 3A). Furthermore, an antibody raised against the middle-portion of SAS-4 likewise had the same perimeter as α-tubulin, in line with the fact that the SAS-4 relative CPAP is a microtubule binding protein (Fig 3A, 3C, #9) (40–42). Thus, α-tubulin and SAS-4 can be used inter-changeably as invariant references in this analysis.

**Figure 3:**
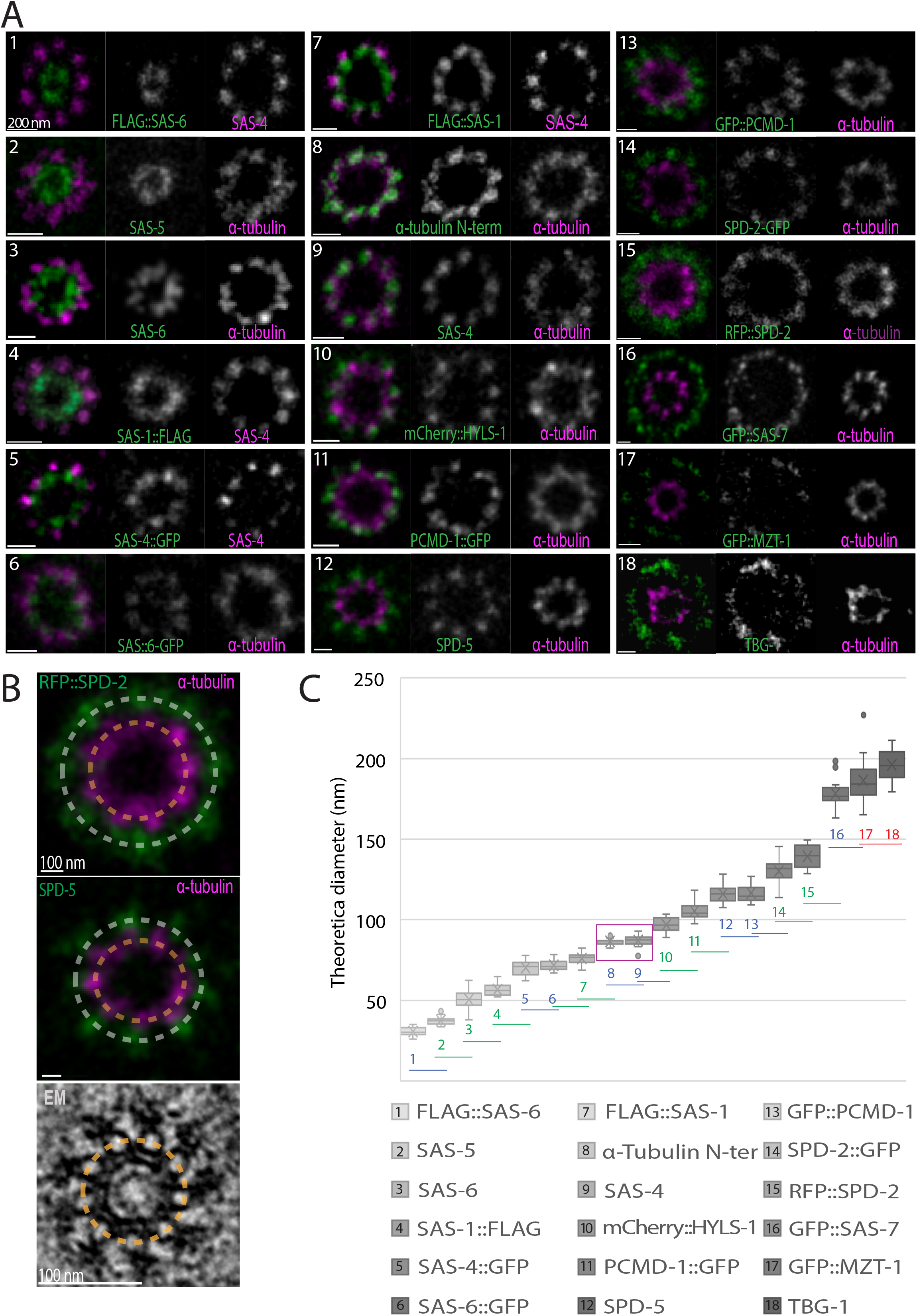
Relative position of centriolar components within the centriole. (A) U-Ex-STED of centrioles from early meiotic prophase stained for the indicated proteins. Each component (green) was imaged together with either α-tubulin (visualized with an antibody recognizing the C-terminus of the protein) or SAS-4 (visualized with an antibody raised against amino acid 350-517 of the protein) (both magenta). (B) Examples of fitted rings on fluorescent signal used to calculate the diameter of each component relative to α-tubulin or SAS-4 standards. In each image, the diameter of the centriolar component (in these cases RFP::SPD-2 (top) and SPD-5 (middle)) and that of the α-tubulin signal were measured along the dashed lines. The perimeter of the centriolar component was then divided by that of the α-tubulin signal. To obtain the theoretical diameter of the component before expansion, this value was normalized by the diameter of microtubules in EM images of centrioles (see Fig. 5). (C) Calculated diameter of each centriolar component as determined in (B), arranged from the smallest to the largest. Magenta box highlights α-tubulin (#8) and SAS-4 (#9). Numbers in the graph indicate the identity of the component. Colors indicate whether the diameter is significantly different from zero (red), one (blue), or two (green) neighboring values (Student’s two tailed t-test, significance p < 0.005). The middle lines of the boxplots correspond to the median, the cross represents the mean, the box includes 50% of values (IQR) and the whiskers show the range of values within 1.5*IQR. N= Flag::SAS-6: 21, SAS-5: 25, SAS-6: 22, SAS-1::FLAG: 10, SAS-4::GFP: 21, SAS-6::GFP: 15, FLAG::SAS-1 15, α-tubulin (N-ter): 19, SAS-4: 22, mCherrry::HYLS-1: 20, PCDM-1::GFP: 24, SPD-5: 20, GFP::PCDM-1: 21, SPD-2::GFP: 25, RFP::SPD-2: 20, GFP::SAS-7: 15, GFP::MZT-1: 20 and TBG-1: 20.

To estimate the ring diameter of each component in non-expanded samples, we determined the diameter of the ring formed by the 9 microtubules in a novel EM dataset of early meiotic prophase centrioles to be 87.9 ± 5.7 nm (N=44, see below), and compared this value to the α-tubulin signal diameter determined with U-Ex-STED (Fig. 3B). Moreover, we found that the α-tubulin diameter determined with U-Ex-STED following correction of the expansion factor (5.2) is similar to that measured for microtubules by EM (88 ± 8 nm; N=38). This standardization method enabled us to estimate the actual diameter of the ring distribution of each protein, going from the smallest one, SAS-6[N], to the largest ones, SAS-7[N], MZT-1 and TBG-1 (Fig. 3C). This analysis established that most components that were shown previously through biochemical and cell biological assays to physically interact are indeed located in close vicinity to one another. This is the case for SAS-6 and SAS-5 (43), SAS-4 and HYLS-1 (17), SAS-7 and SPD-2 (8, 44), SPD-2 and SPD-5 (45), as well as PCMD-1 and SAS-4 or SPD-5 (46).

Overall, U-Ex-STED enabled us to localize in a comprehensive manner centriolar and core PCM component with unprecedented spatial precision.

### 9-fold symmetrical and chirality establishing components of the *C. elegans* centriole

We next addressed whether the ring-like distribution of each centriolar and core PCM component exhibits 9-fold radial symmetry. To this end, we conducted an analysis of the U-Ex-STED data set for each component that is illustrated in the case of α-tubulin in Figure 4A and 4B. First, a circle was drawn along the ring-like signal and an intensity profile measured along this circular line (Fig. 4A, 4B). In the majority of cases, this yielded 9 clearly distinguishable peaks. In ideal top views, with no or very little tilt of the organelle with respect to the imaging axis, the average distance between signal peaks is consistent with the 40° angle expected from a 9-fold radially symmetric structure (Fig. 4C). Importantly, besides α-tubulin, we found a 9-fold symmetric arrangement for SPD-5, PCDM-1[C], SPD-2[C], HYLS-1[N], SAS-4, SAS-6[C] and SAS-1[N] (Fig. 4 D-K, left two panels, raw).

**Figure 4:**
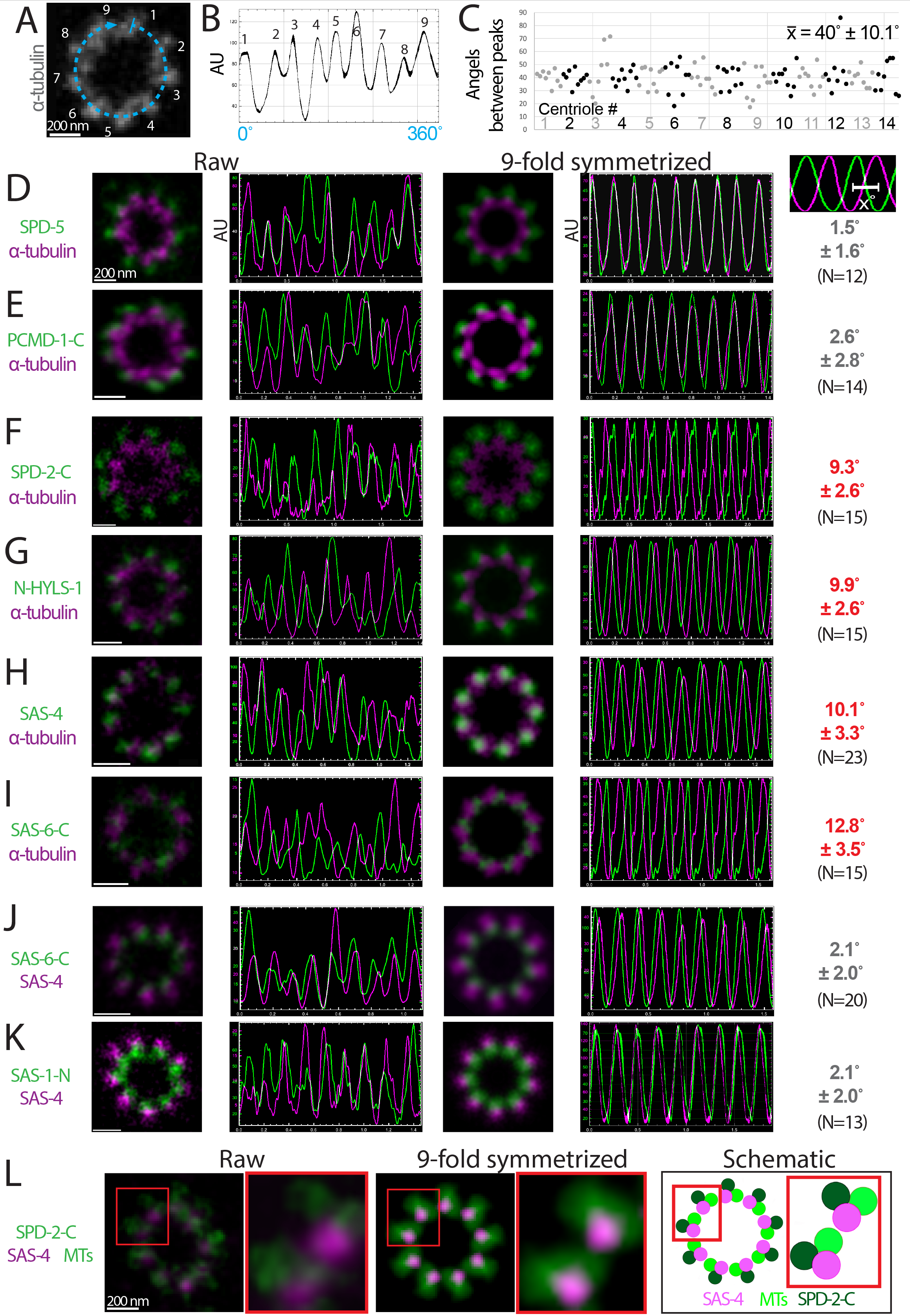
Chiral features of the centriole. (A) U-Ex-STED of a centriole stained for α-tubulin. Numbers correspond to the signal peaks in the intensity profile along the dashed line reported in (B). (B) Signal intensity profile along the dashed line in (A) (4 pixels wide). (C) Angles between peaks of α-tubulin signal intensity profiles in 14 top views of centrioles imaged with U-Ex-STED. Angles were determined by dividing the distance between each neighboring peak by the length of the entire profile, multiplied by 360. Alternating grey and black data points indicated values from each of the 14 centrioles examined. 11/14 centrioles showed 9 clearly discernable signal peaks. 2/14 cases display 8 and 1/14 10 signal peaks. (D-K) U-Ex-STED and plot signal intensity profiles of raw images (left two panels) and corresponding 9-fold symmetrized versions (right) of the indicated pairs of components (green and magenta). In cases top views were slightly tilted, images were circularized before 9-fold symmetrization using the Fiji plugin ‘Transform-Interactive Affine’. The numbers on the very right represent the average offset of the two nearest signal intensity peaks in 9-fold symmetrized images for the two components tested. (L) (Left) U-Ex-STED of centriole from a worm expressing SPD-2::GFP stained for GFP and α-tubulin in the same color (green), as well as for SAS-4 (magenta). (Middle) Corresponding 9-fold symmetrized version. (Right) Schematic representation of the immuno-fluorescence analysis, manually separating the SPD-2::GFP and α-tubulin signals based on their ring diameter. Red boxes are magnified on the right of each image. Note that we have not analyzed HYLS-1 in this manner, as the mCherry::HYLS-1 signal intensity is too weak to this end.

We next investigated whether centriolar proteins thus localized exhibit an offset distribution with respect to microtubules. To that end we examined if the 9-fold radial symmetric distributions are on the same angular axis as the microtubules using the following analysis pipeline. First, we averaged the signals of the microtubules and of the component to be tested by applying 9-fold symmetrization (Fig. S2). Second, we acquired a signal intensity plot along the ring in the resulting symmetrized images for both channels. As expected given the 9-fold symmetrization, in such an analysis individual signal peaks for each channel are ∼40° apart (360° / 9 signal peaks) (Fig 4D-K). Third, signal intensity plots from the two channels are overlaid, and the angular distance between each peak in the α-tubulin channel and the neighboring peak in the second channel determined. In this manner, the average angular offset in each centriole of the component in question versus microtubules is computed.

Strikingly, the above analysis pipeline revealed that components exhibiting a 9-fold symmetric arrangement fall into two well separated groups. In a first two-membered group containing SPD-5 and PCMD-1[C], the offset with respect to microtubules is marginal (<3°) (Fig. 4D, 4E), similar to that of the two antibodies raised against α-tubulin (2.4° ± 1.6, N=10). Therefore, SPD-5 and PCMD-1[C] are not offset with respect to the microtubules. In stark contrast, a second group of components exhibited a clear offset (>9°) with respect to the microtubules, thus leading to a chiral ensemble (Fig. 4F-4K). This second group includes SPD-2[C] and HYLS-1[N], which are both located outside the microtubules (Fig. 4F, 4G). In addition, SAS-4 and SAS-1[N], which both have a ring diameter similar to that of α-tubulin, exhibited an offset with respect to microtubules (Fig. 4H, Fig. 3C). More internally, SAS-6[C] also exhibits a strong offset with respect to microtubules (Fig. 4I). Importantly, we found additionally that SAS-4 and SAS-6[C] are well aligned with one another (Fig. 4J), as are SAS-4 and SAS-1[N] (Fig. 4K). Thus, the offset of SAS-4 exhibits the same handedness with respect to microtubules as that of SAS-6 and SAS-1.

How does the offset handedness of components located outside the microtubules relate to that of those located more centrally? To address this question, we set out to simultaneously examine offset in the angular axis with respect to microtubules of the outer components SPD-2[C] and the inner offset component SAS-4. Because our microscopy setup does not lend itself to performing high-quality 3-color STED, we marked SPD-2[C] and α-tubulin in the same color in this experiment, since they exhibit clearly distinct ring diameters (see Fig. 3A, 3C, # 8 and #14). Importantly, this analysis uncovered that SPD-2[C] and SAS-4 are invariably located on the same side of the microtubules (Fig. 4L). Note that this is regardless of whether SPD-2[C] and SAS-4 are to the left or to the right of the α-tubulin signal, which is expected to depend on whether a centriole is viewed from the proximal or the distal end. We conclude that the offset of the outer component SPD-2[C] and the inner component SAS-4 has the same handedness with respect to microtubules.

### Ultrastructural map of the *C. elegans* gonad centriole

Having achieved precise protein localizations in the gonad centriole with U-Ex-STED, we set out to determine whether we can assign specific centriolar components to specific compartments of the organelle. Given that the ultrastructure of the worm centriole has been best studied in the early embryo (6–8), and considering that the centriole may exhibit tissue-specific features, we set out to directly characterize the ultrastructure of the gonad centriole. Using Correlative Light Electron Microscopy (CLEM) of chemically fixed samples to ease spotting of G2 early meiotic prophase centrioles followed by 50 nm serial sectioning, we acquired the largest EM data set of worm centrioles to date (N=44). As shown in Figure 5A and 5B, we found that peripheral paddlewheels, microtubules, central tube and inner tube are all clearly discernable, as they are in the early embryo (6–8). However, side views established that the centriole is shorter in the gonad than in the early embryo (96.6 ± 7.2 nm as compared to ∼175 nm (7, 8)). Moreover, top view revealed that the paddlewheels are slightly smaller as well (8). Apart from these two differences, we conclude that the overall ultrastructure of the centriole is conserved between the embryo and the gonad.

**Figure 5:**
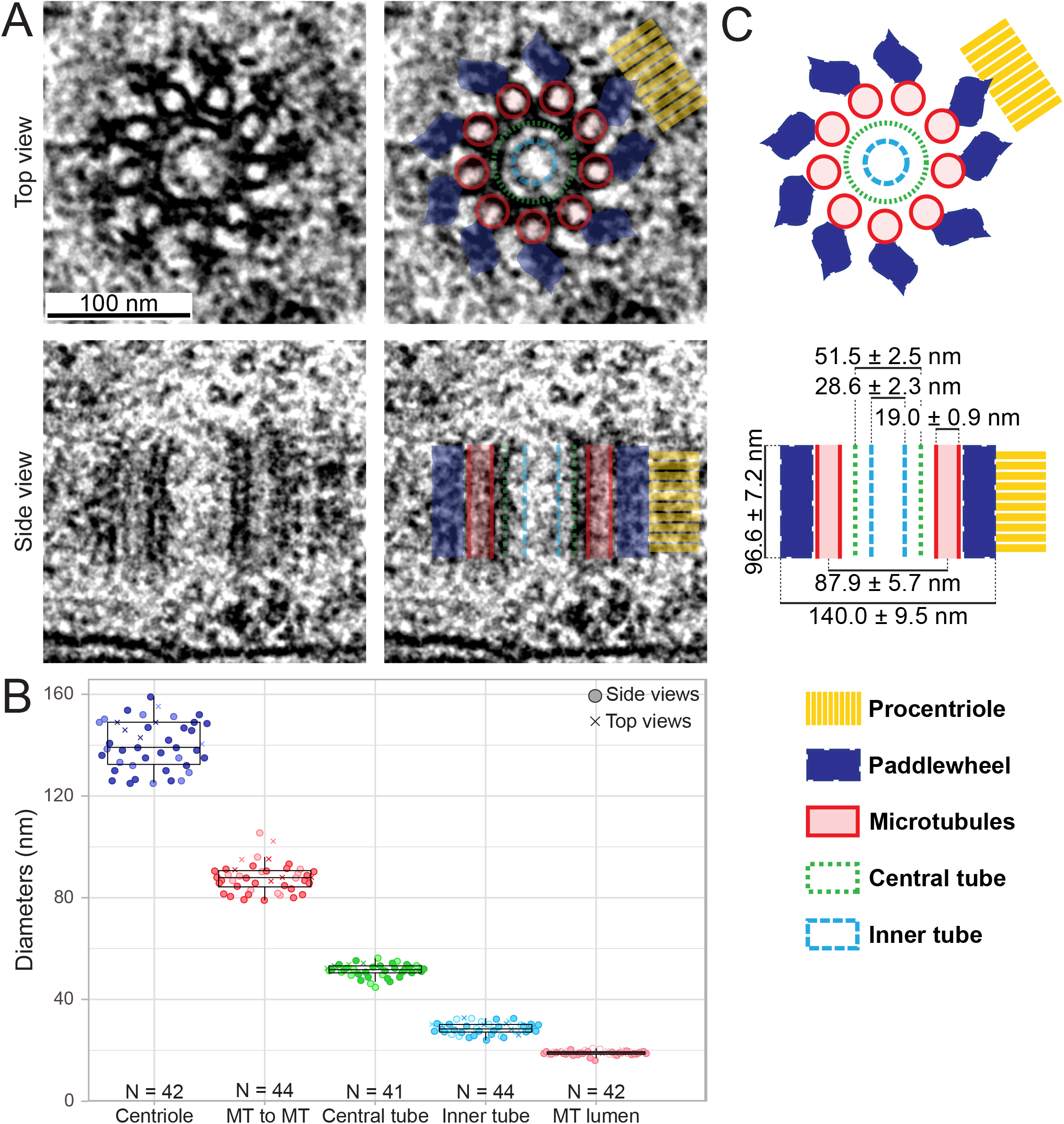
EM analysis of the centriole in the gonad. (A) (Left) EM of top and side views, as indicated, of early meiotic prophase centrioles. (Right) Overlay with distinct ultrastructural compartments as described in (C). (B) Diameters of ultrastructural compartments of the gonad centriole. Top view (crosses) and side views (circles) were analyzed; light and dark shade of colors represent data points from two independent samples. The middle lines of the boxplots correspond to the median, the box includes 50% of values (IQR) and the whiskers show the range of values within 1.5*IQR. (C) Schematic representation of top and side views of centrioles with ultrastructural compartments depicted in colors, as well as average measurements ± SD (see also C). N=38 for centriole height.

Using both top and side views, we determined the diameter of the entire organelle, encompassing the most peripheral paddlewheel features, to be 140.0 ± 9.5 nm (Fig. 5B, 5C). Moreover, the diameters of central tube and inner tube are 51.5 ± 2.5 nm and 28.6 ± 2.3 nm i, respectively, whereas that of the radial array of the microtubules is 87.9 ± 5.7 nm (Fig. 5C). These measurements are in line with those from high-pressure frozen early embryonic centrioles (7, 8), indicating that there is no or marginal shrinkage due to chemical fixation.

The paddlewheels of the centriole in the embryo were reported to exhibit a clockwise twist with respect to the microtubules when viewed from the distal end, using the presence of the procentriole to define the proximal end of the centriole (8). In the gonad, however, where the centriole is shorter, the procentriole often covers the centriole side throughout its entire height (Fig. S3A, top row, N=15). In those cases where the centriole was longer than the cross-sectional diameter of the procentriole, the latter could emanate from either the middle of the centriole or from the vicinity of one of the two ends (Fig. S3B, N=17). Thus, chirality of the centriole cannot be assessed reliably with respect to procentriole orientation in the gonad. Nevertheless, the fact that the procentriole can also emanate from the middle of the centriole raises the possibility that centriole chirality might not be fixed with respect to procentriole orientation.

### Establishing the molecular architecture of the *C. elegans* centriole: beyond microtubules

We set out to determine which centriolar and core PCM proteins correspond to which ultrastructural compartment of the organelle. To better understand the cellular context in which the centriole resides, we conducted tomographic analysis of the EM sections (ET), which revealed a ribosome free area ∼262 ± 26 nm in diameter extending beyond the paddlewheels (Fig. 6A, N=3). This diameter is ∼60 nm larger than that of the largest ring-like distribution observed in this work (see Fig. 3C), raising the possibility that other proteins may be present in this area.

**Figure 6:**
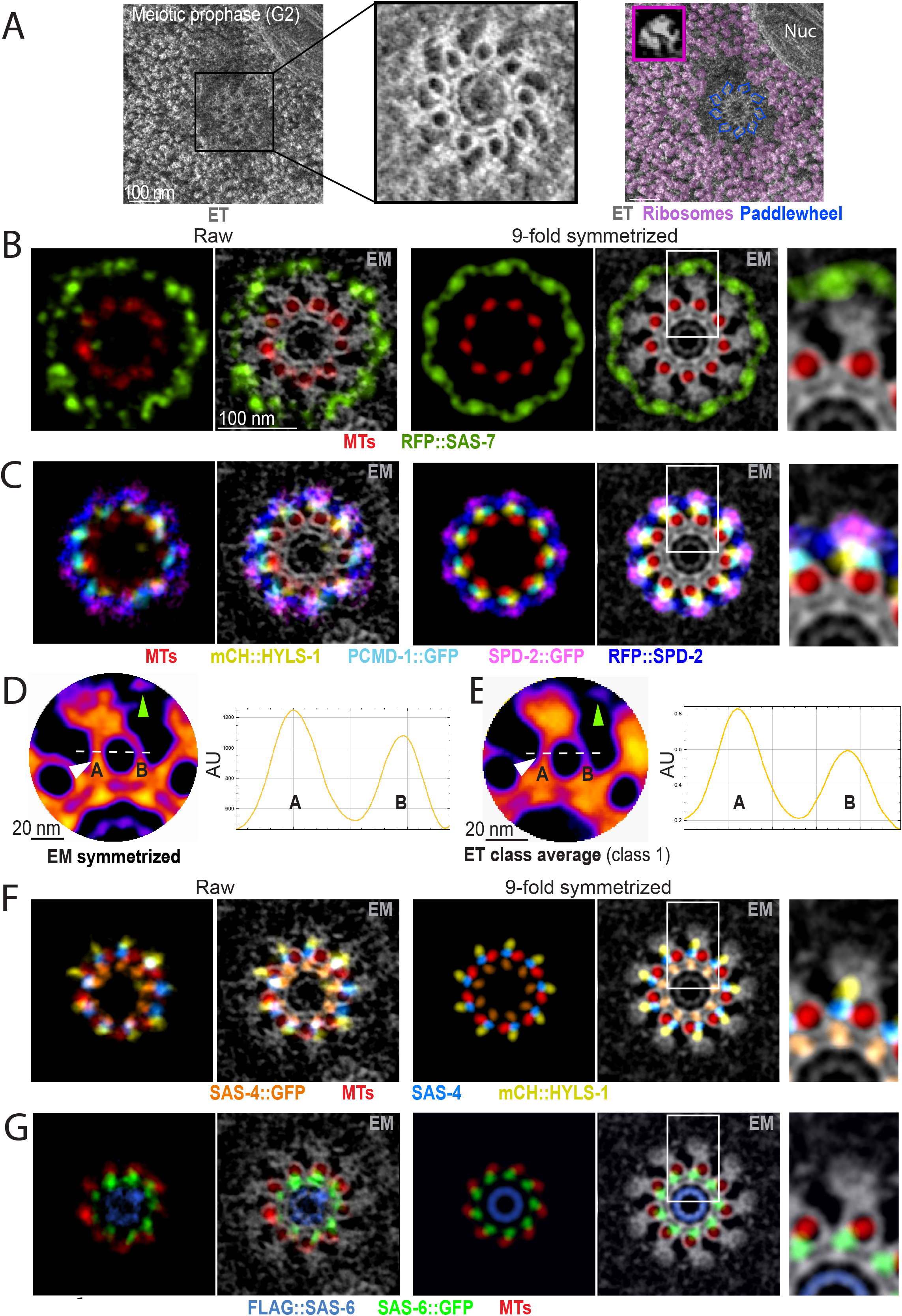
Overlay of EM and U-Ex images. (A) (Left) Max intensity Z-projection of ET of an early meiotic prophase centriole and surrounding region. (Middle) Magnification of the black box in the image on the left. (Right) Manually annotated ribosomes are shown in magenta and paddlewheel structures with dark-yellow outlines. Note that the ribosome-free area extends beyond the paddlewheels. Purple inset shows a magnified ribosome from the same image. (B, C, F, G) Overlay of U-Ex-STED and EM images (inverted grey levels) of centrioles from early meiotic prophase. Circularized images (left two panels), corresponding 9-fold symmetrized versions (next two panels), and magnification of the insets highlighted by the white box (very right). (B) Note that SAS-7 extends beyond the paddlewheel. (C) Overlay of paddlewheel components. (F) Overlay of components around microtubules. (G) Overlay of SAS-6 (N-and C-ter) and microtubules. (D, E) (Left) Magnification of a 9-fold symmetrized centriole imaged by EM (D) and highest populated class from class-averaging of particles containing microtubules and paddlewheels from individual ET tilt series of four centrioles (E) (see Fig. S4). Images are colorized with the LUT “Fire” (low intensities in blue, high intensities in magenta and red). Light green arrowheads point to the small density next to the paddlewheel (IPD), filled white arrowheads to the novel density spanning from the central tube to one side of the microtubule (SCD). (Right) Intensity profiles were obtained along the dashed lines indicated in the images (10 pixels wide). Microtubules displayed consistently more density on the side located under the paddlewheel (marked with an A) compared to the other side (marked with a B).

To map proteins onto ultrastructural compartments, we devised a method that relies on overlaying U-Ex-STED and EM images, using microtubules as a joint registration standard. In brief, we circularized, rotated and size-adjusted jointly the two U-Ex-STED channel signals, aligning the α-tubulin signal with the microtubules in the EM images (Fig. S4). We applied this method initially on the symmetrized images and then likewise adjusted the raw data (Fig. S4). We report the results of this analysis hereafter, starting with the outside of the organelle.

Overlaying the U-Ex-STED and EM data revealed that SAS-7[N] localizes just outside the paddlewheel, partially filling the region devoid of ribosomes surrounding the centriole (Fig. 6B). Four components were found to localize to the paddlewheel: HYLS-1[N], SPD-2, SPD-5 and PCMD-1. SPD-5 and PCMD-1[C] are on the same angular axis as microtubules in the U-Ex-STED data set (see Fig. 4D, 4E), and we indeed find PCMD-1[C] just outwards of microtubules in the overlay, constituting the base of the paddlewheel (Fig. 6C). HYLS-1[N] also localizes to the base of the paddlewheel, but in contrast to SPD-5 and PCMD-1[C], it does so with an offset with respect to the microtubules (Fig. 6C). SPD-2 is the outer-most component of the paddlewheel with the two ends showing distinct distributions: SPD-2[C] appears as foci positioned just outside of microtubules, with an angular offset with respect to them (Fig. 6C, see also Fig. 4F), whereas SPD-2[N] localizes slightly further to the outside as an epitrochoid with 9 lobes extending left and right over the paddlewheel (Fig. 6C). Interestingly, we detected a previously unnoticed small electron-dense region in the EM and ET datasets (see below) located between neighboring paddlewheels (Fig. 6D, 6E, green arrowheads), which can be partially matched with the position of SPD-2[N] in these overlays. We name this density Inter <u>P</u>addlewheel <u>D</u>ensity (IPD). Overall, this analysis reveals in exquisite detail the molecular architecture of components located outside the centriolar microtubules.

### Molecular architecture at the level of the microtubules

We next report the analysis of components located more centrally. Upon careful analysis of the symmetrized EM dataset, we noticed another novel density, which starts from the central tube (Fig. 6D, dashed arrowhead), extends towards and along each microtubule, rendering one side of the microtubule more pronounced than the other (Fig. 6D, white arrowhead). This density displays the same angular offset with respect to the microtubules as the paddlewheel. Since microtubules are not always perfectly perpendicular to the plane of sectioning, we performed ET to obtain *bona fide* top views of microtubules, and thus better analyze this novel density. From individual tilt series of four centrioles, we picked 628 particles containing microtubules and paddlewheels; class-averaging resulted in three well-defined classes containing 92% of input particles (Fig. S5A). In all three classes, the novel density is present on the side of the microtubule above which the paddlewheel is located (Fig. 6E, Fig. S5B). Given that SAS-6, SAS-4 and SAS-1 all display the same angular offset direction with respect to microtubules as the paddlewheel component SPD-2[C] (see Fig. 4F and 4H-L), we propose that these three proteins together could compose this novel offset density. Therefore, we name this novel density “<u>S</u>AS-6/4/1 <u>C</u>ontaining <u>D</u>ensity” (SCD). Overlays of the corresponding U-Ex-STED and EM images indeed revealed perfect alignment of SAS-4 and SAS-6[C] with the SCD, below one side of the microtubule (Fig. 6F and 6G). Moreover, SAS-4[C] overlaps almost perfectly with the SCD at the level of the central tube, and SAS-1[N] has an indistinguishable diameter from SAS-6[C] (see Fig. 3C). Taken together, our data suggest that SAS-4, SAS-6 and SAS-1 form the newly described chiral SCD, with SAS-4 potentially bridging it to HYLS-1.

### The N-terminus of SAS-6 is present at the inner tube and does not form a spiral

We capitalized on the unprecedented high resolution afforded by U-Ex-STED to address whether *C. elegans* SAS-6 forms a ring or instead a steep spiral *in vivo*, as has been hypothesized based on structural and biophysical data (16). The spiral model predicts that SAS-6[N] should be apparent in top views as a small ring with a diameter of ∼4.5 nm (16). Given the ∼14 nm effective lateral resolution achieved using U-Ex-STED, this would appear as a single focus. Contrary to this prediction, we found that the diameter of the ring formed by SAS-6[N] is ∼31 ± 3 nm, overlapping with the inner tube in EM images (Fig. 6G). We noted also that SAS-6[C] localizes ∼41 ± 4 nm away from SAS-6[N], in line with the fact that the coiled-coil domain of SAS-6 is ∼35 nm long and followed by an intrinsically disordered region of ∼90 amino acids (47). Taken together, our observations indicate that, rather than a steep spiral, *in vivo, C. elegans* SAS-6 forms a ring-containing cartwheel.

## Discussion

We deciphered the molecular architecture of the minute *C. elegans* centriole in unprecedented detail by combining U-Ex-STED with EM, thereby localizing twelve centriolar and core PCM proteins to distinct ultrastructural compartments (Fig. 7).

**Figure 7:**
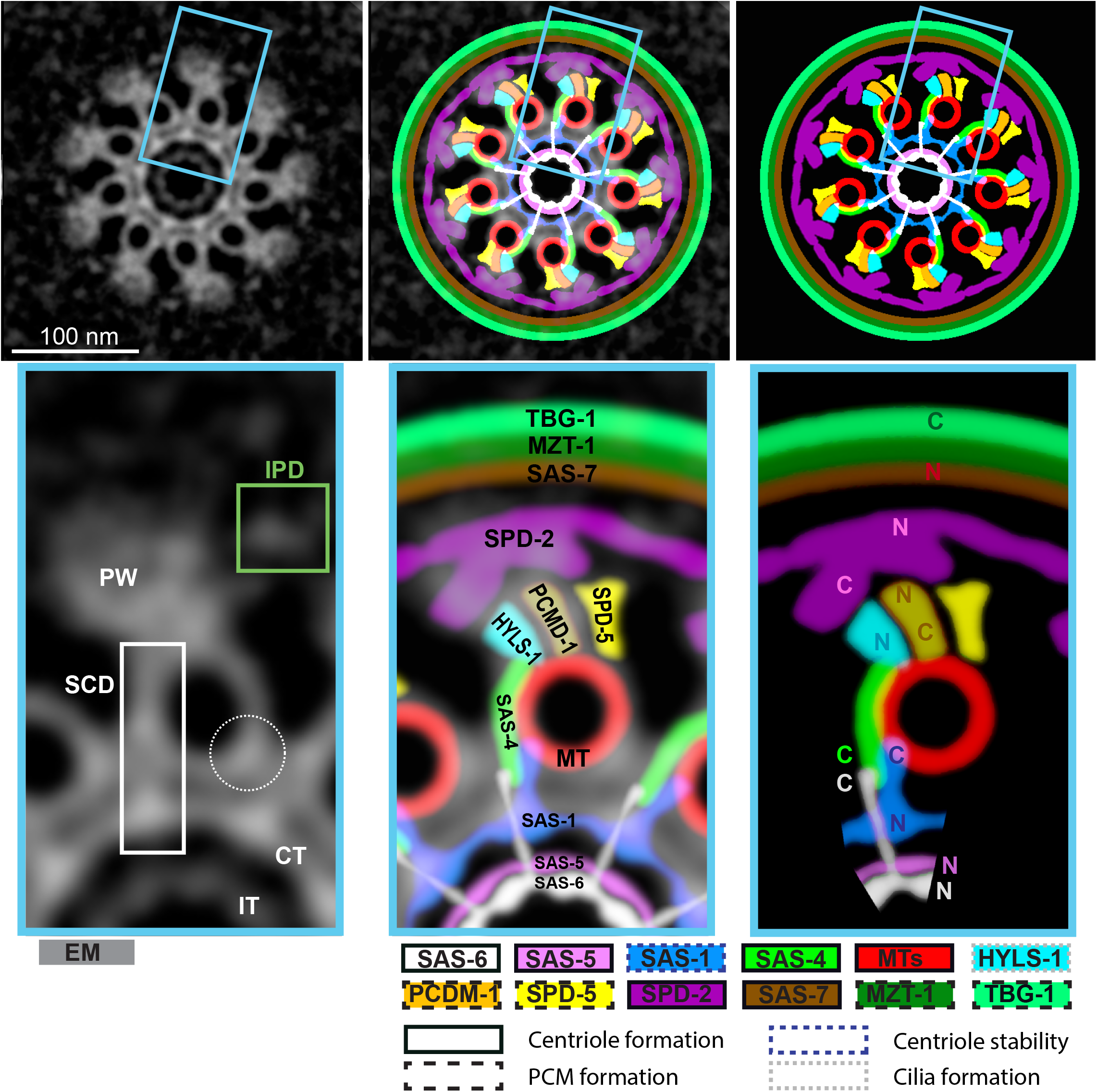
Schematic representation of the localization of components within the centriole. (Top left) 9-fold symmetrized EM image of a centriole. (Top middle) Overlay of the image on the left with the schematic representation on the right. (Top right) Coarse grained schematic representation of the localization within the centriole of the components analyzed in this study. (Bottom left) Magnification of the blue boxed region from the image above. Green box highlights the IPD, white box the SCD. Dashed circle highlights a small density to which no known centriolar protein has been assigned (see Discussion). (Bottom middle and right) Magnifications of the blue boxed regions from the images above, with an indication of the localization of each protein, including where the N-and C-terminus maps when known. No mention of termini is indicated when merely antibodies were utilized.

The precise localization of proteins achieved herein is by and large compatible with, and extends, previous findings. Thus, components that were shown previously through biochemical and cell biological assays to physically interact are indeed located in close vicinity of one another in our map, including SAS-6 and SAS-5 (43), SAS-4 and HYLS-1 (17), SAS-7 and SPD-2 (8, 44), SPD-2 and SPD-5 (45), as well as PCMD-1 with SAS-4 and SPD-5 (46). Other distributions were not necessarily anticipated from prior work. For instance, we found that SAS-7 localizes partly outside the paddlewheel structure, within the zone of ribosome exclusion. SPD-2 and PCMD-1 are not needed for SAS-7 localization, whereas SAS-7 is needed for normal centriolar levels of SPD-2 and PCMD-1, as well as for the integrity of the paddlewheels themselves (8, 46). The localization of SAS-7 outside of SPD-2 and PCMD-1 raises the possibility that SAS-7 functions through a shielding mechanism rather than by recruiting SPD-2 and PCMD-1. Another functionally suggestive distribution uncovered here is that of SAS-1: SAS-1[C] localizes just inside the microtubule ring, in line with the fact that this part of the protein associates with and stabilizes microtubules when ectopically expressed in human cells (18). Therefore, it is tempting to speculate that *C. elegans* SAS-1 maintains centriole integrity by locally exerting a microtubule stabilizing function. The only reported interaction not recapitulated here is that of microtubules with SAS-5, which is mediated by its N-terminus and result in co-localization with the microtubule network upon transfection in COS-7 cells (48). We used antibodies raised against this region of SAS-5, which localize ∼50 nm more centrally than the centriolare microtubules, suggesting that the SAS-5 N-terminus interacts preferentially with another component in the *C. elegans* centriole.

Overlaying EM images with U-Ex-STED images of all known centriolar proteins to date allows us to consider whether there may be centriolar proteins that have not yet been identified in the worm. Such proteins might correspond to EM densities that cannot be readily accounted by the distribution of the proteins assessed here by U-Ex-STED. One such density, distinct from the ICP and the SCD, is apparent on the inner side of microtubules, opposite the SCD (see Fig. 7, dashed circle). This is reminiscent of the region to which Cep135 and Cep97 localize in the fly centriole (34). It has been suggested that a divergent Cep135 protein localizes to centrioles of *C. elegans* during certain developmental stages (49), and it will be interesting to use U-Ex-STED to address whether it maps to this density. Alternatively, it is possible that a segment of the proteins tested here and localizing to such a density would not be apparent from tagging the N-or C-terminal part of the protein, or with some of the antibodies.

Our approach enabled us to probe the higher order oligomerization mechanisms of SAS-6 *in vivo*. Previous structural and biophysical experiments led to the suggestion that such oligomers form a steep spiral instead of a ring as in other systems (16). The steep spiral model predicts that SAS-6[N] appears in top views as a small ring with a diameter of ∼4.5 nm (16). We found instead that SAS-6[N] forms a ring ∼31 ± 3 nm in diameter, which neatly overlays with the inner tube ultrastructural compartment in EM images. We conclude that SAS-6 does not form a steep spiral in the worm and propose instead that the protein forms a ring-containing cartwheel as in other organisms. Alternatively, *C. elegans* SAS-6 may assemble into a shallow spiral. Moreover, we found SAS-6[C] to be positioned ∼41 ± 4 nm away from SAS-6[N], compatible with a cartwheel structure in which the SAS-6 coil-coil domains form spokes extending towards the peripheral microtubules, as in other systems (14, 15). It will be interesting to uncover how the intrinsic properties of *C. elegans* SAS-6 that enable it to form a steep spiral *in vitro* are modulated in the organismal context to adopt a ring-like configuration. This might be aided by interacting proteins such as SAS-5 (44, 48, 50), or by a connection of SAS-6[C] to microtubules. Alternatively, properties of the centriole surface from which the procentriole assembles might impose a different conformation, since the presence of a surface can help constrain the inherent helical properties of *Chlamydomonas reinhardtii* SAS-6 polymers into a ring (51).

Our analysis uncovered offset protein distributions with respect to microtubules, thereby resulting in a chiral centriolar ensemble. Such an offset pertains notably to SAS-6[C] and SAS-4, which coincide with the newly identified SCD, an EM density found centrally and laterally to the microtubule wall. Interestingly, the SCD displays an angular offset with respect to the microtubule in the same direction as the paddlewheel and its constituent SPD-2[C]. Angular offsets with respect to centriolar microtubules occur in other systems. For example, EM studies of centrioles in *Trichonympha* and *Chlamydomonas* uncovered that the pinhead component connecting cartwheel and microtubules exhibits an angular offset with respect to the A microtubule that is on the side of the B and C microtubules (52). Moreover, super-resolution microscopy in Drosophila revealed that the centriolar proteins Cep135 and Ana1 exhibit an angular offset with respect to the A microtubule that is on the other side than the B microtubule (34).

Chirality of the centriole is a signature feature of the organelle observed across the eukaryotic domain of life. However, the potential evolutionary pressure leading to conservation of such chirality is not clear, although an appealing possibility is that this could be optimal for ciliary and flagellar motility. Regardless, it has been suggested that centriolar chirality may be imparted by inherent chiral features of SAS-6 proteins, with chirality in the inner part of the organelle dictating that of more peripheral elements, including microtubules (51). Alternatively, chirality could stem from the fact that microtubules of the procentriole grow with a fixed orientation from the surface of the centriole, with the plus end leading. Therefore, the surface for molecular interaction available on the left and the right side of a microtubule is inherently different. As a result, a protein that interacts with a specific surface on the microtubule wall and that has a fixed orientation along the polymer, such as SAS-4 (see Fig. 7), would necessarily render the centriole chiral. Regardless, it will be important to uncover how chirality of the centriole is established and what its role might be in centriolar biogenesis and function.

It will be of interest to apply the methods developed herein to probe potential variations in the molecular architecture of centrioles in distinct developmental contexts in *C. elegans*. For example, we uncovered here that centriole length in the gonad and the early embryo differ substantially; such a height difference may be accompanied by alterations in molecular architecture. Moreover, these methods can be deployed to interrogate with utmost precision the molecular architecture of centrioles in mutant worms in this genetically tractable organism to further unravel mechanisms of organelle biogenesis and function. Beyond *C. elegans*, such an analytical framework is anticipated to likewise reveal the distribution of centriolar proteins in other systems, and thereby identify conserved and variable features of organelle architecture.

## Supplementary figure legends

Supplementary Figure 1: **Procentriole composition and maturation.**

(A) Widefield image of two pairs of centriole/procentriole in the gonad after U-Ex, stained for SAS-6 and α-tubulin. Note that procentrioles harbor SAS-6 but no α-tubulin at this stage.

(B) U-Ex-STED images of centrioles from meiotic prophase (left) and the mitotic zone (middle and right). Images illustrate that the amount of ZYG-1 on the centriole (but not on the procentriole) varies: during meiotic prophase, ZYG-1 levels on the centriole are very low. For quantification, a line was drawn from the center of the centriole to the outside of the procentriole and the intensity profile along this line measured, as represented by the dashed arrows. (Left) In 95% of meiotic prophase centrioles (19/20), a single ZYG-1 signal intensity peak was detected outside of the centriole peak, directly under the procentriole. (Middle and right) In the mitotic zone, by contrast, whereas 12/25 centrioles exhibited a similar distribution, 13/25 displayed the ZYG-1 signal more prominently than during meiotic prophase, with two clear ZYG-1 peaks, one at the base of the procentriole and one in the middle of the centriolar SAS-6::GFP signal.

(C) 3D SIM sum intensity Z-projected image of a nucleus in the mitotic zone of an expanded gonad. Phosphorylated Histone 3 marks nuclei in mitosis. Insets on the right show that all four centrioles contain α-tubulin, unlike in S or G2 phase.

Supplementary Figure 2: **Schematic of 9-fold symmetrization process.**

Images of centrioles seen from the top were centered in a square ROI. Images were cropped and iteratively rotated by 40° (left). The resulting nine images were arranged in a stack and sum intensity projected (middle). The resulting image represents a 9-fold symmetrized image (right).

Supplementary Figure 3: **Procentriole position along mother centrioles.**

(A, B) (Left) EM side views with centered procentriole (top) and off centered procentriole (bottom) in early meiotic prophase. (Middle) Overlay of schematics with EM images. (Right) Corresponding schematic representations. Paddlewheels are highlighted in blue, microtubules in red and procentrioles in yellow. Dashed lines indicate the middle of the height of centrioles (blue) and procentrioles (yellow).

Supplementary Figure 4: **Schematic of U-Ex and EM images image overlay.**

In EM and U-Ex-STED top views, centrioles with slight tilted orientations were circularized with the Fiji plugin ‘Transform-Interactive Affine’. The grey levels of the EM image were inverted, and the circularized EM and U-Ex-STED images then 9-fold symmetrized as illustrated in Supplementary Figure 2. The perimeters of the microtubule wall in symmetrized EM images and of the α-tubulin signal in symmetrized U-Ex-STED images were measured, and the symmetrized U-Ex-STED image adjusted in dimensions so that the α-tubulin signal had the same perimeter as the microtubule wall in symmetrized EM. To overlay symmetrized images, U-Ex-STED images were rotated so that individual α-tubulin signals perfectly overlap with individual microtubule signals in symmetrized EM images. Thereafter, images were overlayed in individual color channels. The rotational angles and size adjustments applied for symmetrized images were then applied also to the raw non-symmetrized images (indicated by the dashed arrows), which were then treated likewise.

Supplementary Figure 5: **ET class averaging of individual microtubules reveals novel densities.**

(A) 628 particles containing microtubules and paddlewheels were picked from individual ET tilt series of four centrioles. Class-averaging resulted in 12 classes, three of which were well-defined and together contained 92% of input particles (classes 1-3, colorized with the LUT “Fire”, low intensities in blue, high intensities in magenta and red).

(B) Line intensity profile of classes 1-3 along the lines indicated in (A) (10 pixels wide). The microtubule displayed consistently more density on the side located under the paddlewheel (“A”) than on the other side (“B”).

**Table S1.**
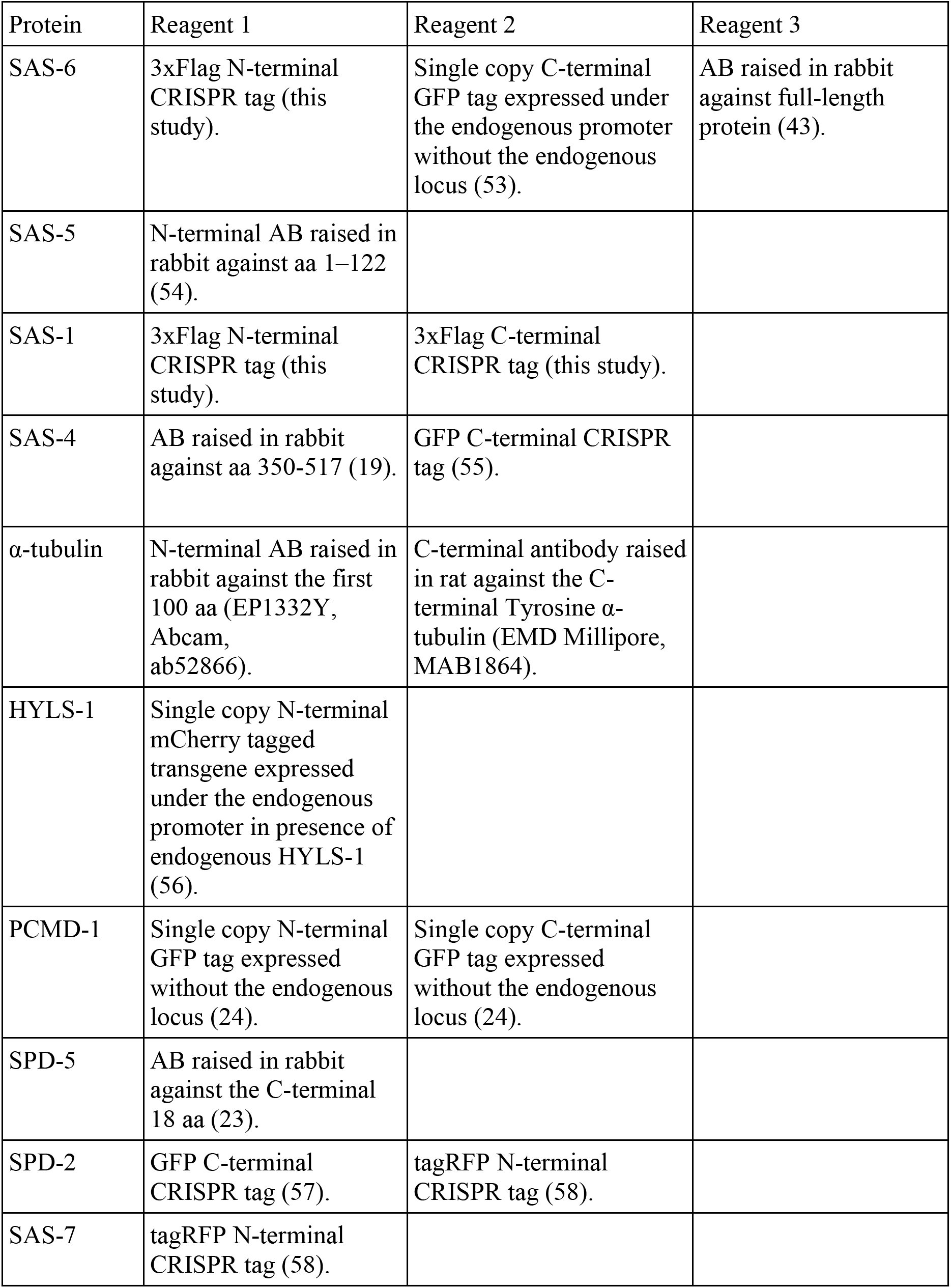
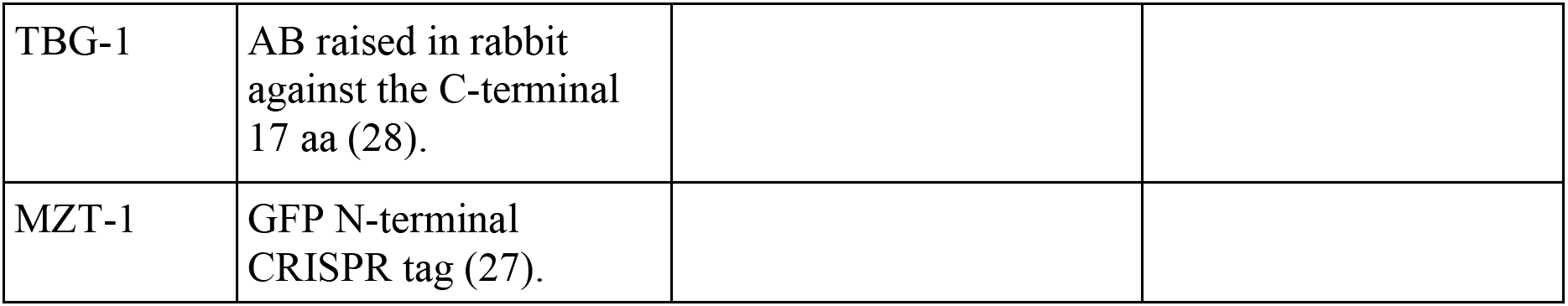
List of Reagents to detect centriolar proteins.

## Materials and Methods

### *C. elegans* culture conditions

Worms were grown on *E. coli* (OP50) seeded NG agar plates at 20°C and age matched as L1 larvae by bleaching gravid adults according to (59). Worms were harvested for ethanol fixation or gonad spreading 24 to 36 hours post L4 stage by washing them off the plate with PBS-T (PBS supplied with 0.1% Tween-20).

### CRISPR/Cas9 genome editing

3xFLAG tagging of SAS-1 and SAS-6 was performed by CRISPR/Cas9 as described in (60). crRNAs were designed using the GUIDE DESIGN tool (http://crispr.mit.edu). Briefly, young adult worms were injected with CRISPR/Cas9 ribonuclear complexes and *dpy-10* was used as the co-injection marker. F1 progenies with roller or dumpy phenotypes were selected and the edits assessed using PCR in the F2 generation, followed by verification by Sanger sequencing. The crRNA sequences were:

SAS-1 N-terminus: ACAATTACTGGTGCCCTTCT(CGG)

SAS-1 C-terminus: CGGATTTGGAGAATATGATG(AGG)

SAS-6 N-terminus: AATTTTGCTAGTCATTTTTG(TGG)

### Ethanol fixation

Worms were washed twice in PBS-T and kept for 30-60 minutes in PBS-T in a 1.5 mL tube to allow emptying of intestines. PBS-T was removed, and 1.5 ml of 100% ethanol then added. Worms were precipitated by gravitation, and the ethanol then removed completely before resuspension of the worms in 25 μL MVD (50% M9, 50% Vectashield (Vector), 0.7 ug/L Hoechst (bisBenzimide H 33258)). Fixed worms were pipetted onto a slide and a 20×40 mm ethanol-washed coverslip was applied with slight pressure.

### Gonad spreading

Spreading of *C. elegans* gonads was performed in a similar manner as in (39). Gonads of ∼ 1000 adult worms were dissected in 30 μL dissection solution (0.2 x PBS, 1:1000 Tween 20) on an ethanol-washed 22×40 mm coverslip. 5-10 μL of dissected gonads were then pipetted onto a new ethanol-washed 22×40 mm coverslip and 50 μL of spreading solution (for one coverslip, 50 μL: 32 μL of Fixative (4% w/v Paraformaldehyde and 3.2% w/v Sucrose in water), 16 μL of Lipsol solution (1% v/v/ Lipsol in water), 2 μL of Sarcosyl solution (1% w/v of Sarcosyl in water)) was added, and gonads were immediately distributed over the coverslip using a pipette tip. Coverslips were left to dry at room temperature followed by incubation at 37°C for 1 hour. Coverslips were either processed for staining and expansion or stored at −80°C.

### Immuno-fluorescence

Dried coverslips were incubated for 20 min in methanol at −20°C. After washing 3 times for 5 min in PBS-T (1x PBS, 1:1000 Tween 20), coverslips were blocked for 20 min in 3% w/v BSA in PBS-T at room temperature. Primary antibody incubation was done overnight at room temperature in a moist chamber at 4°C with primary antibodies diluted in 3% w/v BSA in PBS-T supplemented with 0.05% w/v NaN3. Thereafter, coverslips were washed 3 times 5 min in PBST prior to incubation with secondary antibodies for 2 h at room temperature. After three 5 min washes in PBS-T, the coverslips were mounted on a slide using approximately 20 μL Vectashield (Vector) and sealed with nail polish.

### Ultrastructure expansion microscopy

Dried coverslips were incubated for 20 min in methanol at −20°C and washed 3 times in PBS-T for 5 minutes, followed by two washes in PBS for 5 minutes each. Coverslips were incubated in a 5 cm Petri dish overnight at room temperature in Acrylamide/Formaldehyde solution (1% Acrylamide and 1% Formaldehyde in PBS) under mild agitation. Thereafter, coverslips were washed 3 times 5 min in PBS. For gelation, coverslips were incubated in 50 µl monomer solution (19% (wt/wt) Sodium Acrylate, 10% (wt/wt) Acrylamide, 0.05% (wt/wt) BIS in PBS) supplemented with 0.5% Tetramethylethylenediamine (TEMED) and 0.5% Amonium Persulfate (APS) on a piece of Parafilm for 1 h at 37°C in a moist chamber in the dark. All subsequent steps were carried out under mild agitation at room temperature unless otherwise stated. Gels were incubated for 15 min in denaturation buffer (200 mM SDS, 200 mM NaCl and 50 mM Tris in distilled water, pH=9) in 5 cm Petri dishes followed by incubation for 1 h on a 95°C hot plate in fresh denaturation buffer for denaturation. Gels were transferred to 15 cm Petri dishes washed with distilled water 5 times for 20 min, followed by incubation in distilled water overnight at 4°C. The expansion factor was estimated by measuring the gel size with a ruler.

### Immuno-fluorescence of expanded gels

After expansion, gels were cut in pieces fitting into a 5 cm Petri dish. Prior to staining, gels were blocked for 1 h in blocking buffer (10mM HEPES (pH=7.4), 3% BSA, 0.1% Tween 20, sodium azide (0.05%)), followed by incubation overnight with primary antibodies diluted in blocking buffer. Gels were washed three times in blocking buffer for 10 min each, before incubation with secondary antibodies diluted in blocking buffer (supplemented with 0.7 ug/L Hoechst) at 37°C in the dark for 3 h. Gels were washed three times in blocking buffer for 10 min before transfer into a 10 cm Petri dish for re-expansion by washing 6 times 20 min in distilled water. For imaging, gels were cut and mounted on a 60×24 mm coverslip coated with poly-D-lysine (Sigma, # P1024) diluted in water (2 mg/ml) and supported on both longitudinal sides with capillaries attached with superglue. To prevent drying, the edges of the gel were covered with VaLaP (1:1:1 mixture of petroleum:jelly:lanolin:paraffin wax) and the gel was covered with Halocarbon oil 700 for imaging.

### Antibodies used in this study

Primary antibodies raised in rabbit: SAS-6 (1:1000, (43)), SAS-4 (1:800, (19)), SAS-5 (1 :50, (54)), α-tubulin EP1332Y (1:500, Abcam, ab52866), GFP (1:250, a gift from Viesturs Simanis), SPD-5 (1:250, (23)) TBG-1 (1:500, (28)), tagRFP (1:500, Evrogen, AB232), phospho-histone H3 (ser10) (1:300, Merck, 06-570) and mCherry (1:500, Thermo Fisher, PA5-34974).

Primary antibodies raised mouse: GFP (1:100, Merck, MAB 3580) and FLAG (1:500, Themo Fisher, MA1-91878).

Primary antibodies raised in rat: tyrosine α-tubulin (EMD Milpore, MAB1864).

Secondary antibodies (all used aat 1:1000): donkey anti-rabbit conjugated to Alexa Fluor 594 (Abcam, ab150072), donkey anti-mouse conjugated to Alexa Fluor 594 (Abcam, ab150112072), goat anti-rat conjugated to Alexa Fluor 594 (Thermo Fisher, A11007), goat anti-rabbit conjugated to Alexa Fluor 488 (Thermo Fisher, A11034), goat anti-mouse conjugated to Alexa Fluor 488 (Thermo Fisher, A11001), donkey anti-rat conjugated to Alexa Fluor 488 (Invitrogen, A21208) and goat anti-rabbit Alexa Fluor 647 (Thermo Fisher, A10523).

### Imaging

2D-STED images were acquired on a Leica TCS SP8 STED 3X microscope with a 100 x 1.4 NA oil-immersion objective, using 488 nm and 589 nm excitation, and 592 nm and 775 nm pulsed lasers for depletion. 1 pixel Gaussian blur was applied to all images for analysis and display. For display, brightness and contrast was adjusted in the individual channels.

3D-SIM images were acquired on an inverted Nikon Eclipse Ti instrument, with motorized stage and HXP illumination using an APO TIRF 100 x 1.49 NA oil-immersion objective. Image reconstruction was performed with the NIS Elements software and SUM-intensity projected for analysis and display.

### Determination of effective resolution

The resolution of STED images was determined with 589 nm excitation and depletion with the 775 nm pulsed depletion laser in 10 raw images of α-tubulin using the ImageJ plugin “ImageDecorelationAnalysis” (61), which resulted in a resolution estimate of 73.4 (± 7.96) nm. This resolution was divided by the average expansion factor of 5.2, determined by the perimeter of α-tubulin signals in the U-Ex-STED images divided by the perimeter of microtubules in EM images. SDs of all three measurements were summed up as a percentage of each individual measurement (estimation of resolution in the 10 images, measurements for perimeters of α-tubulin and measurements of perimeters of microtubules).

### CLEM analysis

Gonads of genotype *sas-7(or1940[gfp::sas-7])III; glo-1(zu931)X; itIs37[pie-1p::mCherry::H2B*, unc-119(+)] or *ltSi202[pVV103/ pOD1021; Pspd-2::GFP::SPD-5 RNAiresistant;cb-unc-119(+)]II; sas-7(is1[tagRFP::sas-7+loxP])III; glo-1(zu931)X* were dissected in sperm buffer (50mM Hepes pH7.0, 50mM NACL, 25mM KCL, 5mM CaCl2, 1mM MgSO4, 50mM Glucose, 1mg/ml BSA) and transferred on poly-lysine coated MatTek glass bottom dishes. 3D imaging of gonads was performed using a Nikon Ti2-E epifluorescent microscope equipped with an Andor camera Zyla-4.2P-CL10 before and after ∼2 h 30 min fixation at room temperature in 2.2% glutaraldehyde, 0.9% Paraformaldehyde in Cacodylate buffer 0.05M (pH 7.4), 0.09M sucrose and 0.9mM MgCl2. Briefly, specimens were postfixed in 1% osmium tetroxide, 0.8% potassium ferrocyanide in cacodylate buffer (0.1M, pH 7.2), treated with 0.2% Tannic Acid in 0.05M cocadylate buffer (pH 7.0), stained with 1% uranyl acetate in Sodium Acetate (pH 5.2), dehydrated in an alcohol series and embedded in Hard EPON. 50 nm sections were imaged at 23’000x magnification using a TecnaiSpirit (FEI Company) operated at 80 kV and equipped with an Eagle CCD camera (FEI Company). Using relative positioning of centrioles and nuclei in fluorescence images facilitated the search of centrioles and restricted the number of sections to be imaged. Gaussian blur filtering 1.5 was applied on displayed EM images.

Ultrastructural compartments of the centriole were measured manually using Fiji (62). Each data point is the average of four measurements extracted from lines drawn along the height of the feature. In some cases, ultrastructural compartments could not be measured because they were not visualized accurately, or the view was too tilted. Graphs were generated using PlotsofData (63) and SuperPlotsofdata (64). Procentriole positioning relative to the centriole in Fig. S3 was qualitatively assessed on side views, excluding views that are too tilted.

### Electron tomography

Tilt-series from cryo-fixed sections were acquired on a Tecnai F20 operated at 200 kV (Thermo Fischer Scientific using Thermo Scientific Tomography software in continuous tilt scheme from −60° to +60° in 2° steps at −2.5 µm defocus. Data were recorded with a Falcon III DD camera (Thermo Fisher Scientific) in linear mode at 29’000 × magnification, corresponding to a pixel size of 3.49. Particles were picked from individual tilt images and 2D Class averages were calculated using Relion (65), Xmipp (66), and Eman2 within the Scipion3 (67) framework. Tilt series alignment and tomogram reconstruction was done using EMAN 2.9 (68). Tomogram sub-volumes for the detection of ribosome-free area were extracted using Imod 4.9 (69) and maximum intensity project in Fiji (62).

Worm strains used in this study:

- N2 (Bristol)
- TMD101: *pcmd-1(t3421); mikSi6[pmai-2:GFP::C17D12.7] II* (24)
- TMD117: *pcmd-1(t3421); mikSi9[pmai-2:C17D12.7::GFP] II* (24)
- DAM276 : *ltSi40 [pOD1227; Psas-6::sas-6reencoded::GFP; cb unc-119(+)] II; sas-6(ok2554) IV* (53)
- GZ1934: *sas-1(is7[3xflag::sas-1]) III* (this study)
- GZ1966: *sas-1(is6[sas-1::3xflag]) III* (this study)
- OC994: *sas-4(bs195[sas-4::gfp] III* (a gift from Kevin O’Connell)
- DAM307: *vieSi16[pAD390; Phyls1:mcherry::hyls-1; cb unc-119(+)] IV* (56)
- GZ1528: *spd-2(is2[tagRFP::spd-2 +loxP]) I; sas-7(or1940(gfp::sas-7)) III; glo-1(zu931) X* (this study)
- DAM640: *spd-2(vie4[spd-2::gfp +loxP]) I* (57)
- JLF375: *mzt-1(wow51[GFP:MZT-1]) I; zif-1(gk117) III* (27)
- GZ1929: *sas-6(is10[3xflag::sas-6]) IV* (this study)

## Acknowledgements

We are grateful to the laboratories of Alex Dammermann, Jessica Feldman, Bruce Bowerman, Anthony Hyman, Tamara Mikeladze-Dvali, Kevin O’Connell and Viesturs Simanis for their gift of worm strains and antibodies. Some strains were provided by the *Caenorhabditis* Genetics Center (CGC), which is funded by the NIH Office of Research Infrastructure Programs (P40 OD010440). We thank Graham Knott and Marie Croisier (BioEM platform of the School of Life Sciences, EPFL) for TEM work, as well as Tamara Mikeladze-Dvali, Nils Kalbfuss and Georgios Hatzopoulos for constructive comments on the manuscript. This work was supported in part by the European Union (MCSA-IF 588594 to AW and COFUND-EuroPostdoc 588459 to FS), as well as the Swiss National Science Foundation (grant 310030_197749 to PG).

## References

1. J. Azimzadeh, Exploring the evolutionary history of centrosomes. Philos. Trans. R. Soc. Lond. B. Biol. Sci. 369, 20130453 (2014).

2. M. Bornens, The centrosome in cells and organisms. Science 335, 422–426 (2012).

3. M. Winey, E. O’Toole, Centriole structure. Philos. Trans. R. Soc. B Biol. Sci. 369, 20130457 (2014).

4. R. Uzbekov, C. Prigent, Clockwise or anticlockwise? Turning the centriole triplets in the right direction! FEBS Lett. 581, 1251–1254 (2007).

5. D. Ito, M. Bettencourt-Dias, Centrosome Remodelling in Evolution. Cells 7, E71 (2018).

6. E. T. O’Toole, et al., Morphologically distinct microtubule ends in the mitotic centrosome of Caenorhabditis elegans. J. Cell Biol. 163, 451–456 (2003).

7. L. Pelletier, E. O’Toole, A. Schwager, A. A. Hyman, T. Müller-Reichert, Centriole assembly in Caenorhabditis elegans. Nature 444, 619–623 (2006).

8. K. Sugioka, et al., Centriolar SAS-7 acts upstream of SPD-2 to regulate centriole assembly and pericentriolar material formation. eLife 6, e20353 (2017).

9. N. Wolf, D. Hirsh, J. R. McIntosh, Spermatogenesis in males of the free-living nematode, Caenorhabditis elegans. J. Ultrastruct. Res. 63, 155–169 (1978).

10. N. Banterle, P. Gönczy, Centriole Biogenesis: From Identifying the Characters to Understanding the Plot. Annu. Rev. Cell Dev. Biol. 33, 23–49 (2017).

11. E. A. Nigg, A. J. Holland, Once and only once: mechanisms of centriole duplication and their deregulation in disease. Nat. Rev. Mol. Cell Biol. 19, 297–312 (2018).

12. S. Gomes Pereira, M. A. Dias Louro, M. Bettencourt-Dias, Biophysical and Quantitative Principles of Centrosome Biogenesis and Structure. Annu. Rev. Cell Dev. Biol. 37, 43–63 (2021).

13. M. Delattre, C. Canard, P. Gönczy, Sequential protein recruitment in C. elegans centriole formation. Curr. Biol. CB 16, 1844–1849 (2006).

14. I. Vakonakis, The centriolar cartwheel structure: symmetric, stacked, and polarized. Curr. Opin. Struct. Biol. 66, 1–7 (2021).

15. P. Guichard, V. Hamel, P. Gönczy, The Rise of the Cartwheel: Seeding the Centriole Organelle. BioEssays News Rev. Mol. Cell. Dev. Biol. 40, e1700241 (2018).

16. M. Hilbert, et al., Caenorhabditis elegans centriolar protein SAS-6 forms a spiral that is consistent with imparting a ninefold symmetry. Proc. Natl. Acad. Sci. U. S. A. 110, 11373– 11378 (2013).

17. A. Dammermann, et al., The hydrolethalus syndrome protein HYLS-1 links core centriole structure to cilia formation. Genes Dev. 23, 2046–2059 (2009).

18. L. von Tobel, et al., SAS-1 is a C2 domain protein critical for centriole integrity in C. elegans. PLoS Genet. 10, e1004777 (2014).

19. S. Leidel, P. Gönczy, SAS-4 is essential for centrosome duplication in C elegans and is recruited to daughter centrioles once per cell cycle. Dev. Cell 4, 431–439 (2003).

20. L. Pintard, B. Bowerman, Mitotic Cell Division in Caenorhabditis elegans. Genetics 211, 35–73 (2019).

21. C. A. Kemp, K. R. Kopish, P. Zipperlen, J. Ahringer, K. F. O’Connell, Centrosome maturation and duplication in C. elegans require the coiled-coil protein SPD-2. Dev. Cell 6, 511–523 (2004).

22. L. Pelletier, et al., The Caenorhabditis elegans centrosomal protein SPD-2 is required for both pericentriolar material recruitment and centriole duplication. Curr. Biol. CB 14, 863– 873 (2004).

23. D. R. Hamill, A. F. Severson, J. C. Carter, B. Bowerman, Centrosome maturation and mitotic spindle assembly in C. elegans require SPD-5, a protein with multiple coiled-coil domains. Dev. Cell 3, 673–684 (2002).

24. A. C. Erpf, et al., PCMD-1 Organizes Centrosome Matrix Assembly in C. elegans. Curr. Biol. CB 29, 1324–1336.e6 (2019).

25. S. Strome, et al., Spindle dynamics and the role of gamma-tubulin in early Caenorhabditis elegans embryos. Mol. Biol. Cell 12, 1751–1764 (2001).

26. E. Hannak, et al., The kinetically dominant assembly pathway for centrosomal asters in Caenorhabditis elegans is gamma-tubulin dependent. J. Cell Biol. 157, 591–602 (2002).

27. M. D. Sallee, J. C. Zonka, T. D. Skokan, B. C. Raftrey, J. L. Feldman, Tissue-specific degradation of essential centrosome components reveals distinct microtubule populations at microtubule organizing centers. PLoS Biol. 16, e2005189 (2018).

28. E. Hannak, M. Kirkham, A. A. Hyman, K. Oegema, Aurora-A kinase is required for centrosome maturation in Caenorhabditis elegans. J. Cell Biol. 155, 1109–1116 (2001).

29. J. M. Schumacher, N. Ashcroft, P. J. Donovan, A. Golden, A highly conserved centrosomal kinase, AIR-1, is required for accurate cell cycle progression and segregation of developmental factors in Caenorhabditis elegans embryos. Dev. Camb. Engl. 125, 4391– 4402 (1998).

30. M. Srayko, S. Quintin, A. Schwager, A. A. Hyman, Caenorhabditis elegans TAC-1 and ZYG-9 form a complex that is essential for long astral and spindle microtubules. Curr. Biol. CB 13, 1506–1511 (2003).

31. J. M. Bellanger, P. Gönczy, TAC-1 and ZYG-9 form a complex that promotes microtubule assembly in C. elegans embryos. Curr. Biol. CB 13, 1488–1498 (2003).

32. L. Gartenmann, et al., A combined 3D-SIM/SMLM approach allows centriole proteins to be localized with a precision of ∼4-5 nm. Curr. Biol. CB 27, R1054–R1055 (2017).

33. T. T. Yang, et al., Super-resolution architecture of mammalian centriole distal appendages reveals distinct blade and matrix functional components. Nat. Commun. 9, 2023 (2018).

34. Y. Tian, et al., Superresolution characterization of core centriole architecture. J. Cell Biol. 220, e202005103 (2021).

35. D. Gambarotto, et al., Imaging cellular ultrastructures using expansion microscopy (U-ExM). Nat. Methods 16, 71–74 (2019).

36. N. Sahabandu, et al., Expansion microscopy for the analysis of centrioles and cilia. J. Microsc. 276, 145–159 (2019).

37. F. Chen, P. W. Tillberg, E. S. Boyden, Optical imaging. Expansion microscopy. Science 347, 543–548 (2015).

38. P. M. Fox, et al., Cyclin E and CDK-2 regulate proliferative cell fate and cell cycle progression in the C. elegans germline. Dev. Camb. Engl. 138, 2223–2234 (2011).

39. A. Woglar, et al., Quantitative cytogenetics reveals molecular stoichiometry and longitudinal organization of meiotic chromosome axes and loops. PLoS Biol. 18, e3000817 (2020).

40. L.-Y. Hung, H.-L. Chen, C.-W. Chang, B.-R. Li, T. K. Tang, Identification of a novel microtubule-destabilizing motif in CPAP that binds to tubulin heterodimers and inhibits microtubule assembly. Mol. Biol. Cell 15, 2697–2706 (2004).

41. A. Sharma, et al., Centriolar CPAP/SAS-4 Imparts Slow Processive Microtubule Growth. Dev. Cell 37, 362–376 (2016).

42. X. Zheng, et al., Molecular basis for CPAP-tubulin interaction in controlling centriolar and ciliary length. Nat. Commun. 7, 11874 (2016).

43. S. Leidel, M. Delattre, L. Cerutti, K. Baumer, P. Gönczy, SAS-6 defines a protein family required for centrosome duplication in C. elegans and in human cells. Nat. Cell Biol. 7, 115–125 (2005).

44. S. Li, et al., A map of the interactome network of the metazoan C. elegans. Science 303, 540–543 (2004).

45. M. Boxem, et al., A protein domain-based interactome network for C. elegans early embryogenesis. Cell 134, 534–545 (2008).

46. L. Stenzel, et al., PCMD-1 bridges the centrioles and the pericentriolar material scaffold in C. elegans. Dev. Camb. Engl. 148, dev198416 (2021).

47. D. Kitagawa, et al., Structural basis of the 9-fold symmetry of centrioles. Cell 144, 364–375 (2011).

48. S. Bianchi, et al., Interaction between the Caenorhabditis elegans centriolar protein SAS-5 and microtubules facilitates organelle assembly. Mol. Biol. Cell 29, 722–735 (2018).

49. E. Holzer, C. Rumpf-Kienzl, S. Falk, A. Dammermann, A Modified TurboID Approach Identifies Tissue-Specific Centriolar Components In C. elegans. 2021.12.20.473533 (2021).

50. R. Qiao, G. Cabral, M. M. Lettman, A. Dammermann, G. Dong, SAS-6 coiled-coil structure and interaction with SAS-5 suggest a regulatory mechanism in C. elegans centriole assembly. EMBO J. 31, 4334–4347 (2012).

51. N. Banterle, et al., Kinetic and structural roles for the surface in guiding SAS-6 self-assembly to direct centriole architecture. Nat. Commun. 12, 6180 (2021).

52. P. Guichard, et al., Native architecture of the centriole proximal region reveals features underlying its 9-fold radial symmetry. Curr. Biol. CB 23, 1620–1628 (2013).

53. D. Serwas, A. Dammermann, Ultrastructural analysis of Caenorhabditis elegans cilia. Methods Cell Biol. 129, 341–367 (2015).

54. M. Delattre, et al., Centriolar SAS-5 is required for centrosome duplication in C. elegans. Nat. Cell Biol. 6, 656–664 (2004).

55. J. Magescas, J. C. Zonka, J. L. Feldman, A two-step mechanism for the inactivation of microtubule organizing center function at the centrosome. eLife 8, e47867 (2019).

56. C. Schouteden, D. Serwas, M. Palfy, A. Dammermann, The ciliary transition zone functions in cell adhesion but is dispensable for axoneme assembly in C. elegans. J. Cell Biol. 210, 35–44 (2015).

57. T. Laos, G. Cabral, A. Dammermann, Isotropic incorporation of SPD-5 underlies centrosome assembly in C. elegans. Curr. Biol. CB 25, R648–649 (2015).

58. K. Klinkert, et al., Aurora A depletion reveals centrosome-independent polarization mechanism in Caenorhabditis elegans. eLife 8, e44552 (2019).

59. T. Stiernagle, Maintenance of C. elegans. WormBook Online Rev. C Elegans Biol., 1–11 (2006).

60. A. Paix, A. Folkmann, D. Rasoloson, G. Seydoux, High Efficiency, Homology-Directed Genome Editing in Caenorhabditis elegans Using CRISPR-Cas9 Ribonucleoprotein Complexes. Genetics 201, 47–54 (2015).

61. A. Descloux, K. S. Grußmayer, A. Radenovic, Parameter-free image resolution estimation based on decorrelation analysis. Nat. Methods 16, 918–924 (2019).

62. J. Schindelin, et al., Fiji: an open-source platform for biological-image analysis. Nat. Methods 9, 676–682 (2012).

63. M. Postma, J. Goedhart, PlotsOfData-A web app for visualizing data together with their summaries. PLoS Biol. 17, e3000202 (2019).

64. S. J. Lord, K. B. Velle, R. D. Mullins, L. K. Fritz-Laylin, SuperPlots: Communicating reproducibility and variability in cell biology. J. Cell Biol. 219, e202001064 (2020).

65. J. Zivanov, et al., New tools for automated high-resolution cryo-EM structure determination in RELION-3. eLife 7, e42166 (2018).

66. C. O. Sorzano, et al., Semiautomatic, high-throughput, high-resolution protocol for three-dimensional reconstruction of single particles in electron microscopy. Methods Mol. Biol. Clifton NJ 950, 171–193 (2013).

67. J. M. de la Rosa-Trevín, et al., Scipion: A software framework toward integration, reproducibility and validation in 3D electron microscopy. J. Struct. Biol. 195, 93–99 (2016).

68. G. Tang, et al., EMAN2: an extensible image processing suite for electron microscopy. J. Struct. Biol. 157, 38–46 (2007).

69. J. R. Kremer, D. N. Mastronarde, J. R. McIntosh, Computer visualization of three-dimensional image data using IMOD. J. Struct. Biol. 116, 71–76 (1996).

